# The marine gastropod *Conomurex luhuanus* (Strombidae) has high-resolution spatial vision and eyes with complex retinas

**DOI:** 10.1101/2021.12.21.473630

**Authors:** Alison R. Irwin, Suzanne T. Williams, Daniel I. Speiser, Nicholas W. Roberts

## Abstract

All species within the conch snail family Strombidae possess large camera-type eyes that are surprisingly well-developed compared to those found in most other gastropods. Although these eyes are known to be structurally complex, very little research on their visual function has been conducted. Here, we use isoluminant expanding visual stimuli to measure the spatial resolution and contrast sensitivity of a strombid, *Conomurex luhuanus*. Using these stimuli, we show that this species responds to objects as small as 1.06° in its visual field. We also show that *C. luhuanus* responds to Michelson contrasts of 0.07, a low contrast sensitivity between object and background. The defensive withdrawal response elicited by visual stimuli of such small angular size and low contrast suggests that conch snails may use spatial vision for the early detection of potential predators. We support these findings with morphological estimations of spatial resolution of 1.04 ± 0.14°. These anatomical data therefore agree with the behavioural measures and highlight the benefits of integrating morphological and behavioural approaches in animal vision studies. Furthermore, using contemporary imaging techniques including serial block-face scanning electron microscopy (SBF-SEM), in conjunction with transmission electron microscopy (TEM), we found that *C. luhuanus* have more complex retinas, in terms of cell type diversity, than previous studies of the group have discovered using TEM alone. We found the *C. luhuanus* retina is comprised of six cell types, including a newly identified ganglion cell and accessory photoreceptor, rather than the previously described four cell types.

**Summary statement:** Behavioural trials indicate the eyes of conch snail species *Conomurex luhuanus* provide high-resolution spatial vision, and morphological examination reveals the retina contains more cell types than those of other gastropods.

## 1 Introduction

The form and function of eyes vary widely across the animal kingdom, with well-established associations between structure and aspects of performance. Two functional parameters often used to describe the visual performance of eyes are angular resolution (a measure of the smallest object that can be resolved by an eye) and intensity contrast sensitivity (the difference in the perceived brightness that makes an object distinguishable from its background), hereon referred to as ‘contrast sensitivity’ (Land and Nilsson, 2012). As well as functional performance, eyes vary in complexity; generally, more complex organs are comprised of a higher number of components (McShea, 2000; Oakley and Rivera, 2008; Arendt et al., 2009).

One of the most diverse animal groups in terms of visual system complexity are the Mollusca, which reflects the vast range of lifestyles in the group (Messenger, 1981; Serb and Eernisse, 2008). Within the gastropods, eye types vary from simple pits to complex camera eyes (Serb and Eernisse, 2008), yet, despite this diversity, gastropod visual systems and their functional parameters remain relatively unexplored compared to other groups.

Species from the tropical marine family Strombidae may have eyes with the finest spatial resolution of any gastropod. Based on eye morphology, some strombid species are thought to be capable of resolving objects with angular sizes as small as ca. 1° in their visual field (Seyer, 1994). If these estimates are correct, this is surprisingly high acuity vision for a group of slow-moving and herbivorous gastropods. The only other gastropods with similarly high-resolution vision are pelagic sea slugs *(Pterotracheoidea*), which use their eyes to find prey (Land 1982). Furthermore, strombid eyes may be the largest of any non-cephalopod mollusc, reaching up to ca. 2 mm in diameter (Gillary and Gillary, 1979). The eyes of strombids also have a complex architecture: their retinas contain ca. 50,000 tightly packed photoreceptors, at least four cell types (Gillary and Gillary, 1979; Ozaki, 1986), and their spherical lenses appear to have a graded refractive index to minimise spherical aberration (Seyer, 1994). Moreover, electrophysiological investigations into the visual system of strombid species *Conomurex luhuanus* found evidence for multiple light responses within the retina, consistent with the presence of different types of photoreceptors and with some degree of neural processing occurring in the retina (Gillary, 1974, 1977). Together, these morphological (Gillary and Gillary, 1979; Seyer, 1994) and physiological (Gillary, 1974, 1977) studies suggest strombids have complex eyes that provide them with fine spatial resolution and high contrast sensitivity (Seyer 1994). However, these predictions have yet to be verified with behavioural investigations.

In this study, we integrated behavioural and morphological approaches to explore the visual performance and retinal ultrastructure of the strombid, *C. luhuanus*. This species has been the focus of previous morphological studies on eye structure and retinal ultrastructure using traditional histological methods (Gillary and Gillary, 1979; Ozaki et al., 1986), in addition to physiological investigations of the retina (Gillary, 1974, 1977). We revisited this visual system with contemporary serial block-face SEM (SBF-SEM; Denk and Horstmann, 2004) techniques, together with TEM images, to classify cell types in the retina of *C. luhuanus* and discuss their possible functions. Given that functional properties of eyes are closely associated with the visual needs of their bearers (Nilsson, 2013), this combined behavioural and morphological approach of assessing spatial resolution and contrast sensitivity increases our understanding of strombid behaviour and ecology and may help to explain why these gastropods possess larger and more complex eyes than other gastropods.

## 2 Methods

### 2.1 Sample collection

Adult *C. luhuanus* (n=20, shell length 44–52 mm; shell apex to siphonal canal) were purchased from Tropical Marine Centre (TMC), Bristol, UK and held at the University of Bristol, where they were lodged in tanks (39 litres) with seawater at a density of 4–5 conch snails per tank. Seawater in the holding system was maintained at temperature 25–26° C and salinity 1.025–1.027 sg under a filtration system and was partially siphoned and replaced weekly to avoid accumulation of nitrates. Snails grazed on algae on the tank surfaces and within the substrate, supplemented by food pellets (TMC Gamma NutraShots *Calanus*) which were added twice weekly. Aquaria were illuminated with LED lamps under a 12 hr:12 hr light:dark cycle (lights on from 07:00 to 19:00). Experiments commenced one week after the animals’ arrival at the laboratory and were performed within the next three weeks. All experiments were conducted in accordance with the University of Bristol code of ethics for animal experimentation; approval was given by the University Animal Welfare and Ethical Review Body (AWERB) with University Investigation Number UIN/20/006. The sample size of 20 animals was chosen using G*Power v. 3.1 (Heinrich-Heine-Universität), which suggested 19 as an adequate sample size to perform the statistical tests required, with an additional animal for use in SBF-SEM studies.

### 2.2 Visual behaviour experiments

#### 2.2.1 Experimental setup

Animals were held in a glass experimental container, lightly restrained by fabric tape via magnets attached to the ends of the tape and beneath the container. The magnets allowed some freedom of movement whilst keeping the position and orientation of the snails consistent. Seawater from the holding tanks was used to fill the experimental container, with seawater changed between individuals to maintain water temperature and visibility. The experimental container was positioned in the centre of a camera tent, raised above the floor of the tent to attach the magnets (Fig. 1a). One wall of the camera tent was removed and replaced with a liquid-crystal display (LCD) monitor (Fig. 1a) on which a series of visual stimuli (described below) were displayed. This monitor, together with the camera tent, also prevented the animal from responding to slight movements around the laboratory. Animals were positioned 5 cm from the monitor and allowed to acclimatise for 10 minutes, or until they extended their eyestalks and proboscis and began grazing normally, encouraged by food placed inside the experimental tank.

**Figure 1.**
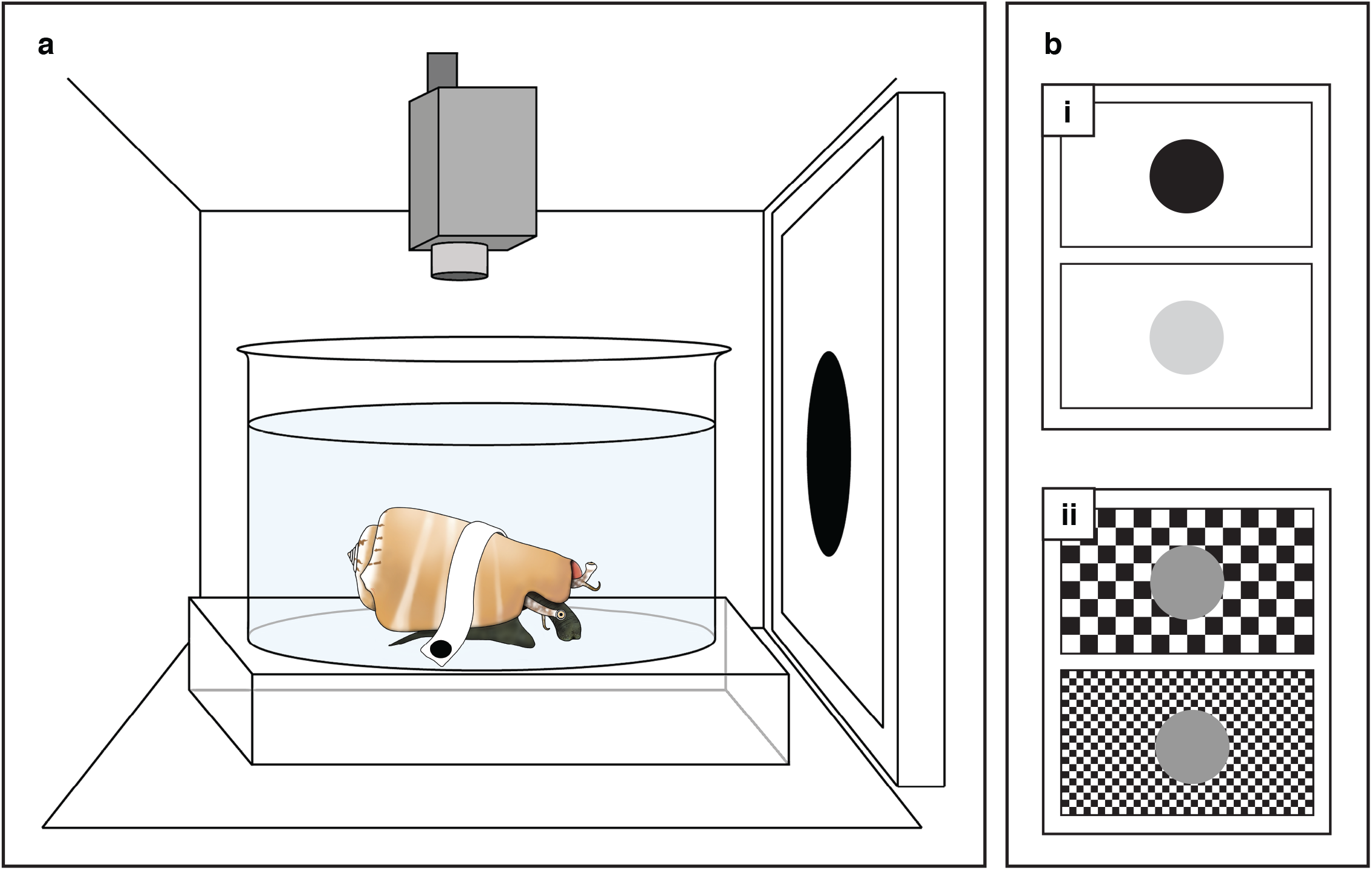
Experimental design. (a) *Conomurex luhuanus* was held in position within the container, placed inside a camera tent. One side of the tent was replaced with an LCD monitor (left), and behaviour was filmed from above (top). (b) Visual stimuli (expanding circles, which expand in the visual field to mimic the approach of a predator; see Movie 1) of varying (i) Michelson contrast and (ii) visual angle were presented on the monitor.

#### 2.2.2 Behavioural assay

Behavioural reactions by conch snails to an expanding visual stimulus (see section 2.3.3 below) were filmed from above with a digital video camera (Canon UK) through a hole in the camera tent (Fig. 1a). Video sequences were synchronised to stimulus events using a single frequency beep produced at the start and end of each stimulus, heard only through headphones. Changes in animal behaviour before, during, and after the stimulus presentations were visually identified from video playback, without knowledge of which stimuli were played in each video during scoring of animal behaviour. Behavioural changes before and during stimulus presentations were divided into seven categories to describe actions concerning the proboscis (1–3) and eyestalks (ommatophores) (4–5): (1) stop feeding; (2) partial proboscis withdrawal; (3) full proboscis withdrawal; (4) partial eyestalk withdrawal; (5) full eyestalk withdrawal (see Table 1 for full descriptions and Movie 1 for video clips of behaviours). The key change in behaviour noted in the period after stimulus presentations was the re-emergence time, defined as the time taken after the maximum response was observed for the eyestalks and the proboscis to re-emerge fully extended from the shell and for the snail to resume normal grazing behaviour.

**Table 1.** Description of behavioural transitions of the *Conomurex luhuanus* eyestalk and proboscis recorded during trials, in sequential order for proboscis and eyestalk, respectively (some transitions featured in Video 1).

|  | Behaviour | Description of behaviour |
| --- | --- | --- |
| Proboscis | Stop feeding | Animal stops feeding, visible when the animal no longer moves its proboscis in a sweeping motion across the aquarium floor |
|  | Partial proboscis withdrawal | Proboscis is retracted towards the body but remains outside the shell. |
|  | Quick, full proboscis withdrawal | Proboscis is retracted inside the shell (not seen by camera). |
| Eyestalk | Partial eyestalk withdrawal | Eyestalks are withdrawn towards the body, are outside or partially inside shell (but remain visible to camera). |
|  | Full eyestalk withdrawal (eyes inside shell, not visible to camera) | Eyestalks are retracted completely inside the shell (eyes not seen by camera). |

#### 2.2.3 Expanding stimulus

Within each experiment (contrast sensitivity and spatial resolution experiments, as outlined below), a series of expanding 2-dimensional stimuli were presented in a randomised order on the monitor screen, with 3-minute intervals between each stimulus presentation (or until conch snails resumed normal grazing behaviour for 3 minutes). Each individual was tested twice for both contrast sensitivity and spatial resolution experiments, with a rest period of at least two days between each of the four tests. Starting with no stimulus present on the screen background, a circular stimulus rapidly expanded (Fig. 1b; Movie 1), simulating, to the eye of the conch snails, the direct approach of a circular object with an angular size that increased from 0° to 83° of the animal’s visual field. Stimuli for the contrast sensitivity and spatial resolution experiments were produced using Matlab v.R2020a (The Math Works, Inc.) or Microsoft PowerPoint (Microsoft Corporation), respectively. Stimuli for each experiment differed with respect to their expansion rates, as described below, so results are not directly comparable between experiments.

##### Behavioural Experiment 1 - Contrast sensitivity experiment

The contrast stimulus was composed of a white background with an expanding circle, which enlarged at an exponential rate over a period of 10s (Fig. 1bi). Variation of the monitor intensity input values for the object produced nine looms with varying differences in intensity between the object and the background (pixel byte values: background = 255; object = 0, 50, 100, 150, 175, 200, 210, 220, 230), reported as Michelson contrasts (parameters close to threshold chosen based on initial observations). A grey loom on an identical background (Michelson contrast 0) was used as the control. The contrast threshold was determined by finding the stimulus with the lowest contrast that elicited a response.

##### Behavioural Experiment 2 - Spatial resolution experiment

The isoluminant spatial resolution stimulus was composed of a black-and-white checkerboard (pixel byte values: black, 0; white, 255) background with a grey expanding circle (pixel byte value: 153). The intensity value of the grey circle was calibrated to match the mean brightness of the white and black squares in the checkerboard. When the stimulus was presented, this circle enlarged at a constant rate over a period of 5s (Fig. 1bii). The size of the squares on the checkerboard background was varied to create eight different stimuli, with the widths of squares ranging in angular size from 0.3 to 3.2°. These sizes were chosen based on initial observations of *C. luhuanus* behaviour and estimates of angular resolution in conch snails from anatomical data (Gillary and Gillary, 1979; Seyer, 1994). If the eyes of conch snails were not able to resolve the black and white squares, the object and the background would appear isoluminant (of equal luminance, i.e., the mid-grey of the object) to the animal, and it should not perceive the first-order motion. Therefore, the visual acuity threshold of conch snails was determined by finding the finest checkboard (checks with smallest angular size) against which animals responded to the isoluminant expanding stimulus. As a measure of spatial resolution, we estimated the minimum resolvable angle (αmin) as twice the angular width of the smallest check to which animals responded.

#### 2.2.4. Statistical analysis of behavioural data

In calculating the probability of an individual showing a behavioural response for each stimulus type, responses where the only behavioural transition observed was ‘stop feeding’ (Table 1) were excluded to reduce the likelihood of a false positive. Wilson score intervals were calculated using the sample size for the experiment and the number of positive responses. We used Spearman’s rank correlation coefficient (SRCC) to investigate whether there was a significant relationship between the response probability and Michelson contrast (contrast sensitivity experiment) or the angular size of the checks in the background (spatial resolution experiment). We also used SRCC to analyse whether there was significant correlation between re-emergence time and Michelson contrast or angular size of checks, and used paired Wilcoxon tests to explore whether there was a significant difference between the median response probability of the two repeats for each of the experiments.

We further analysed the results of these trials by using Fisher’s exact test (FET) to compare the number of individuals that responded to each loom to the number that responded to the control stimulus. To account for multiple comparisons in the contrast sensitivity experiment (nine treatments and one control), we applied a Bonferroni correction.

### 2.3 SBF-SEM

#### 2.3.1 Specimen fixation and embedding

One eye from a specimen of *C. luhuanus* was prepared for SBF-SEM work according to the following protocol. Prior to dissection, the specimen was anaesthetized in a saturated solution of 7.3% MgCl_2_.6H_2_0 mixed with filtered seawater for 30 minutes, at which point the eyestalk withdrawal reflex was absent (Gillary and Gillary, 1979). The right eye, along with anterior parts of the eyestalk, was removed and the animal allowed to recover in a seawater tank. The sample was fixed in a 4% PFA 2.5% glutaraldehyde fixative in 0.1M sodium cacodylate buffer pH 7.3 for 24 hrs on ice with gentle rotation to enable diffusion of the fixatives, and stored in a fresh batch of fixative at 4° C.

To prepare the sample for sectioning, a razor blade was used to slice down the centre of the eye along the sagittal plane, just off the midline (in order to section as close to the middle of the eye as possible), and one half was placed within an automated specimen preparation set-up (Leica EM TP, Leica Biosystems UK) which ran the following steps based on the protocol of the National Center for Microscopy and Imaging Research, University of California, San Diego, CA (Deerinck et al., 2010). The tissue was washed in cold sodium cacodylate buffer (5 × 5 minutes) and fixed with a fresh solution of 1.5% potassium ferrocyanide and 2% aqueous osmium tetroxide in 0.1M cacodylate buffer for 1 hour, at 4°C. At room temperature (RT) the sample was washed with diH_2_O (5 × 5 minutes), incubated with 1% thiocarbohydrazide for 20 minutes, and again washed with diH_2_O (5 × 5 minutes). Then, the sample was fixed in 2% osmium tetroxide in diH_2_0 for 45 minutes, and subsequently washed with diH2O at RT (5 × 5 minutes). The sample was stained/fixed in 1% uranyl acetate (aqueous) overnight at 4° C and subsequently washed at RT with diH_2_O (5 × 5 minutes), and then with 0.03 M aspartic acid pH 5.5 (2 × 10 minutes). The sample was stained with Walton’s lead aspartate pH 5.5 at 60° C for 30 minutes and washed with 0.03 M aspartic acid pH 5.5 (2× 10 minutes), and then diH_2_O (5 × 5 minutes), at RT. An ethanol dehydration series followed: 30%, 50%, 70% (at 4° C for 10 minutes each), 90% (at RT for 10 minutes), 100% (anhydrous; at RT for 4 × 15 minutes), propylene oxide (2 × 15 minutes). The sample was left in 1:1 propylene oxide: hard Durcupan^™^ mix (HDM; Sigma-Aldrich) for 1.5 hours, then in 100% HDM overnight, and finally in fresh HDM for 2 × 3 hours. Tissue was then embedded in a silicon rubber mould and polymerised at 60° C for 48 hours.

#### 2.3.2 Specimen mounting and SBF-SEM imaging

Conventional unstained sections were cut from the fixed and embedded eye with an ultramicrotome (Reichert Ultracut S). These sections were examined with a Tecnai T12 transmission electron microscope (Thermo Fisher Scientific UK) to obtain high-resolution TEM images of the retina and ascertain the quality of fixation prior to SBF-SEM. The resin-embedded tissue was then mounted on an aluminium specimen pin (Gatan, Pleasanton, CA) using silver epoxy glue. The resin block was further trimmed with a glass knife to 1.0 mm × 1.0 mm so tissue in the matrix was exposed on all four sides, with any excess silver epoxy trimmed from around the embedded tissue. The entire surface of the specimen was sputter-coated with a thin layer of gold/palladium. The block was aligned to a 3View microtome (Gatan, USA) mounted in a Zeiss Gemini SEM 450, and a 100 nm thin sectioning of the surface begun. The block surface was imaged using BSE mode (backscattered electron detector, Gatan Onpoint detector, Gatan USA) over an area of 204.8 mm by 204.8 mm in the x,y, at a resolution of 50 nm per raw pixel. The full SBF-SEM run removed 100 sections (100 nm thick), with the block face imaged after each removal.

#### 2.3.3 Annotation and volume segmentation of retinal cells

To analyse retinal cells in a three-dimensional (3D) reconstruction, we traced a single cell through serial semi-thin sections, similar to procedures used to show visual structures in mice and sea spiders (Mustafi et al., 2011; Lehmann et al., 2012; Helmstaedter et al., 2013). Volume Graphics VGStudio Max v. 2.2 was used to segment retinal cells and their nuclei, pigment granules, filaments and phagosomes in 3D reconstruction. Reconstructed cells were used to measure the mean cell volume and mean total volume of pigment granules per cell via VGStudio Max, from which the density of pigment within each cell type was calculated. Counts of each cell type were made in the nuclear layer using Fiji (Schindelin et al., 2012), with cells included in the count if the cell nucleus, a key identifiable characteristic of each cell type, was visible. Estimates of the total number of each cell type per eye were calculated from cell counts within the area of retina sectioned (2.048 × 10**-**_3_ mm^2^) and the total area of the retina measured previously for the same species (1.7 mm^2^; Gillary and Gillary, 1979), rounded to the nearest 100 cells.

#### 2.3.4 Estimates of sensitivity and spatial resolution

The anatomical data was used to estimate the angular resolution of the eyes of *C. luhuanus*. Angular resolution was calculated as twice that of the inter-receptor angle (Δ*ϕ*; Land and Nilsson, 2012; Eqn 1), using the following formula:

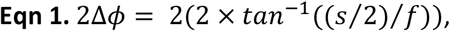

where *s* is the separation of the rhabdom centres and *f* is the focal length. Focal length was estimated as *f* = 2 × *r* (where *r* is the radius of the lens) based on previous measurements of focal length in conch snails (Seyer, 1994). The absolute sensitivity of the eye under a standard luminance (*S*) was also calculated via the same method used to estimate sensitivity in the eye of strombid *Lobatus raninus* (Seyer, 1994), in addition to other animal groups (Kirschfeld, 1974; Land, 1981; Land and Nilsson, 2012; Eqn 2), using the following formula:

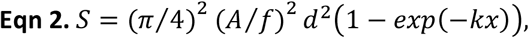

where *A* is the aperture of the eye, *d* is the rhabdom diameter, and *x* is the rhabdom length. Lastly, *k* is the absorption coefficient of the photoreceptors, 0.0067 mm^-1^, the measured value in lobster rhabdoms (Bruno et al., 1977) which was also used by Seyer (1994). While *S* is not directly comparable with contrast sensitivity behaviour experiments, this estimated value is nevertheless a useful metric in discussions of eye function.

## 3 Results

### 3.1 Behavioural experiments

#### 3.1.1 Expanding visual stimuli elicit defensive behavioural responses from conch snails

*C. luhuanus* responded to expanding visual stimuli with a series of defensive behaviours involving retraction of the ommatophores and proboscis (Table 1). These responses are distinguishable from normal grazing activity where the ommatophores and proboscis are extended, the latter moving constantly in a searching motion to feed (Movie 1). These behavioural responses (Fig. 2) are more easily separable into a sequence when the visual stimuli expand more slowly (Fig. 2a), instead of when rapidly expanding stimuli were used (Fig. 2b). Over the expansion period of the stimulus, the following behaviours were observed, in sequential order: stop feeding; proboscis and eyestalks partially withdrawn towards the body (remaining outside the shell; Table 1; Fig. 2a; Movie 1); eyestalks and proboscis fully retracted inside the shell and no longer able to be seen by the camera (Table 1; Fig. 2a; Movie 1).

**Figure 2.**
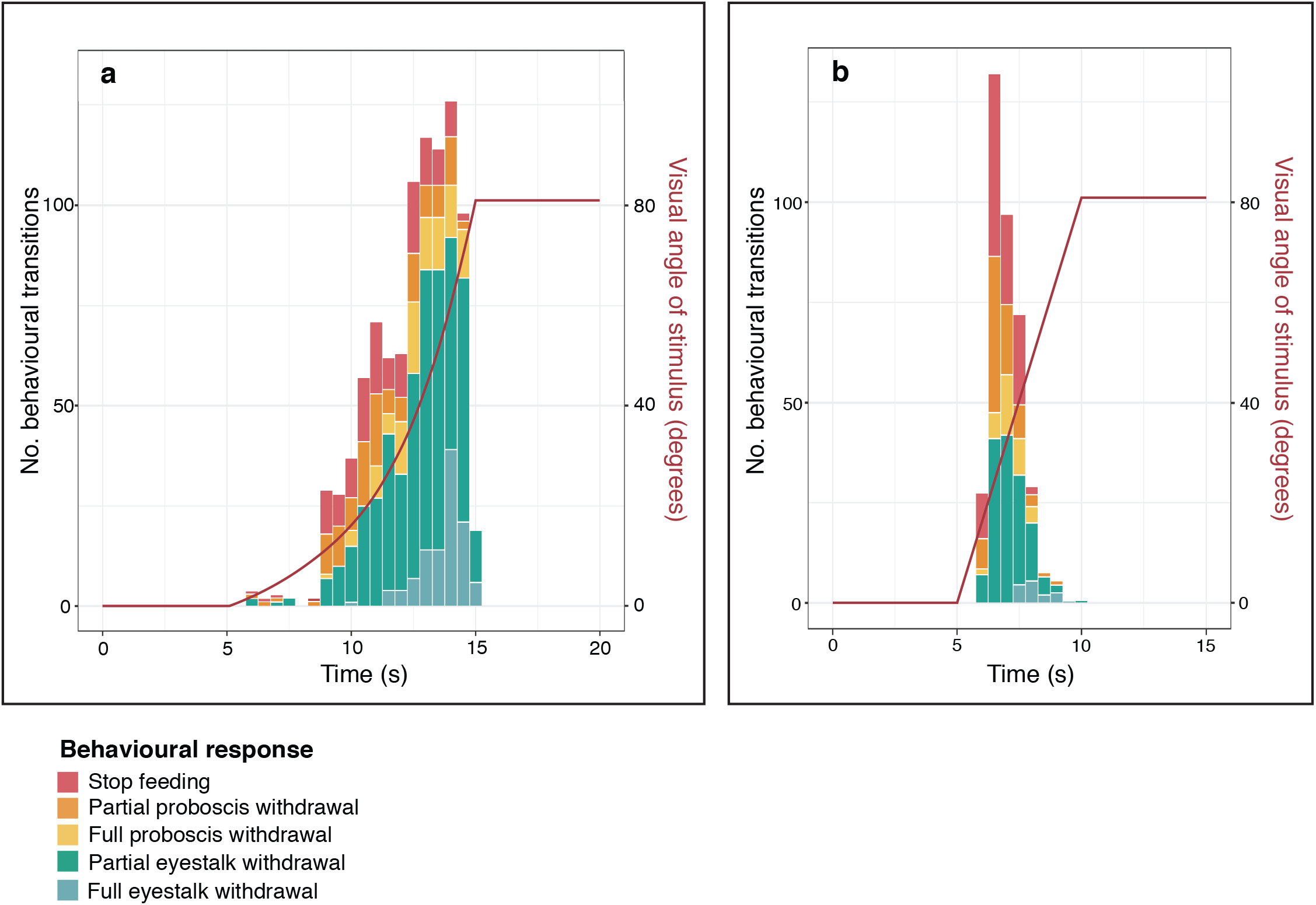
Total number of each *Conomurex luhuanus* behavioural transition (left y-axis; see Table 1 for full behaviour descriptions) displayed over the course of expanding stimulus presentations, plotted for both expanding stimulus types: (a) slow stimulus (contrast sensitivity experiment, n = 20) and (b) fast stimulus (spatial resolution experiment, n = 19). Dark red lines represent the angular size of the stimuli (right y-axis, log scale), which increases (a) exponentially or (b) linearly with time. Each experiment included two replicates for every animal used.

Defensive behaviour exhibited by the conch snails increased considerably during the rapid expansion phase of the expanding circle; after the stimulus had subtended 37.9° of the visual field, 62% of total behavioural transitions in this experiment and 92% of full eyestalk withdrawals were recorded (Fig. 2a). Few behavioural responses (10% of total behavioural transitions in this experiment) occurred when the stimulus was 11.8–16.8° in angular size, and only one animal fully withdrew its eyes before the stimulus reached 27.5° in the visual field (Fig. 2a). Fewest responses (0.01% of total behavioural transitions in this experiment) were recorded during the slow expansion phase of the expanding circle, when the stimulus had subtended 2.3–9.8° of the visual field (Fig. 2a). These were only initial changes in behaviour (stop feeding, partial withdrawal of proboscis and eyestalk towards the shell; Table 1; Movie 1); no full withdrawal responses occurred during this period.

#### 3.1.2 *C. luhuanus* responds to Michelson contrasts of 0.07

The probability of a conch snail showing a behavioural response to a looming visual stimulus is positively correlated with the magnitude of the Michelson contrast of the stimulus (SRCC, *ρ* = 0.942, n = 40, p < 0.001; Fig. 3). In this experiment, 20% of individuals responded to looming visual stimuli with a Michelson contrast of 0.07 (FET, n = 40, p = 0.002), and 62.5% of individuals responded to those with a Michelson contrast of 0.10 (FET, n = 40, p < 0.001) (Fig. 3). No individuals responded to looming stimuli with a Michelson contrast of 0.05 (FET, n = 40, p = 1). The re-emergence time also showed a strong positive correlation with Michelson contrast magnitude (SRCC, *ρ* = 0.833, n = 40, p = 0.015; Fig. 3). There was no significant difference in the median response probability between the two repeats of this experiment (Wilcoxon, *V* = 12, n = 9, p = 0.281).

**Figure 3.**
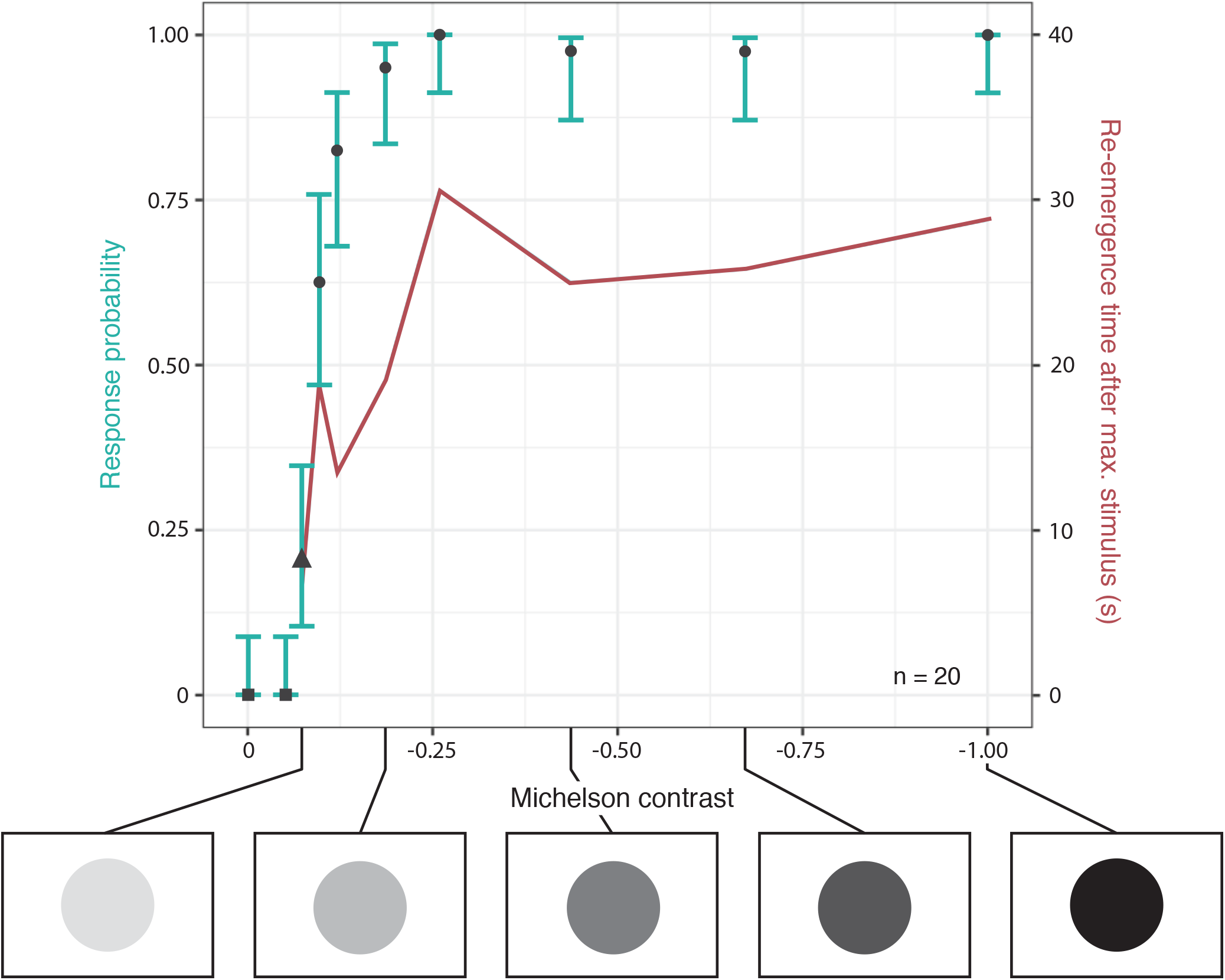
Response probabilities (black shapes) of *Conomurex luhuanus* to expanding stimuli varying in Michelson contrast (below, not to scale): square, p = 1; triangle, p = <0.05; circle, p = <0.01 following Bonferroni correction (n = 20). Error bars (green) are Wilson score intervals. Re-emergence time (red) is the time taken post-stimulus for ommatophores to return to their extended position and normal grazing to resume. This experiment included two replicates for every animal used.

#### 3.1.3 *C. luhuanus* has a spatial resolution of 1.06 degrees

Animals responded to an expanding isoluminant stimulus against a black and white checkerboard pattern consisting of checks with angular widths of 0.53° (FET, n = 38, p < 0.001) and above (Fig. 4a). From this response, the minimum resolvable angle of *C. luhuanus* was 1.06°, twice the angular width of the square checks. No animals responded to an expanding isoluminant stimulus against a black and white checkerboard pattern consisting of checks with angular widths of 0.40° (FET, n = 38, p = 1.000; Fig. 4a). The probability of an individual showing a behavioural response was positively correlated with the angular sizes of the checks in the checkerboard background (SRCC, *ρ* = 0.928, n = 38, p < 0.001; Fig. 4a). There was no significant difference in the median response probability between the two repeats of this experiment (Wilcoxon, *V* = 4, n = 8, p = 0.419).

**Figure 4.**
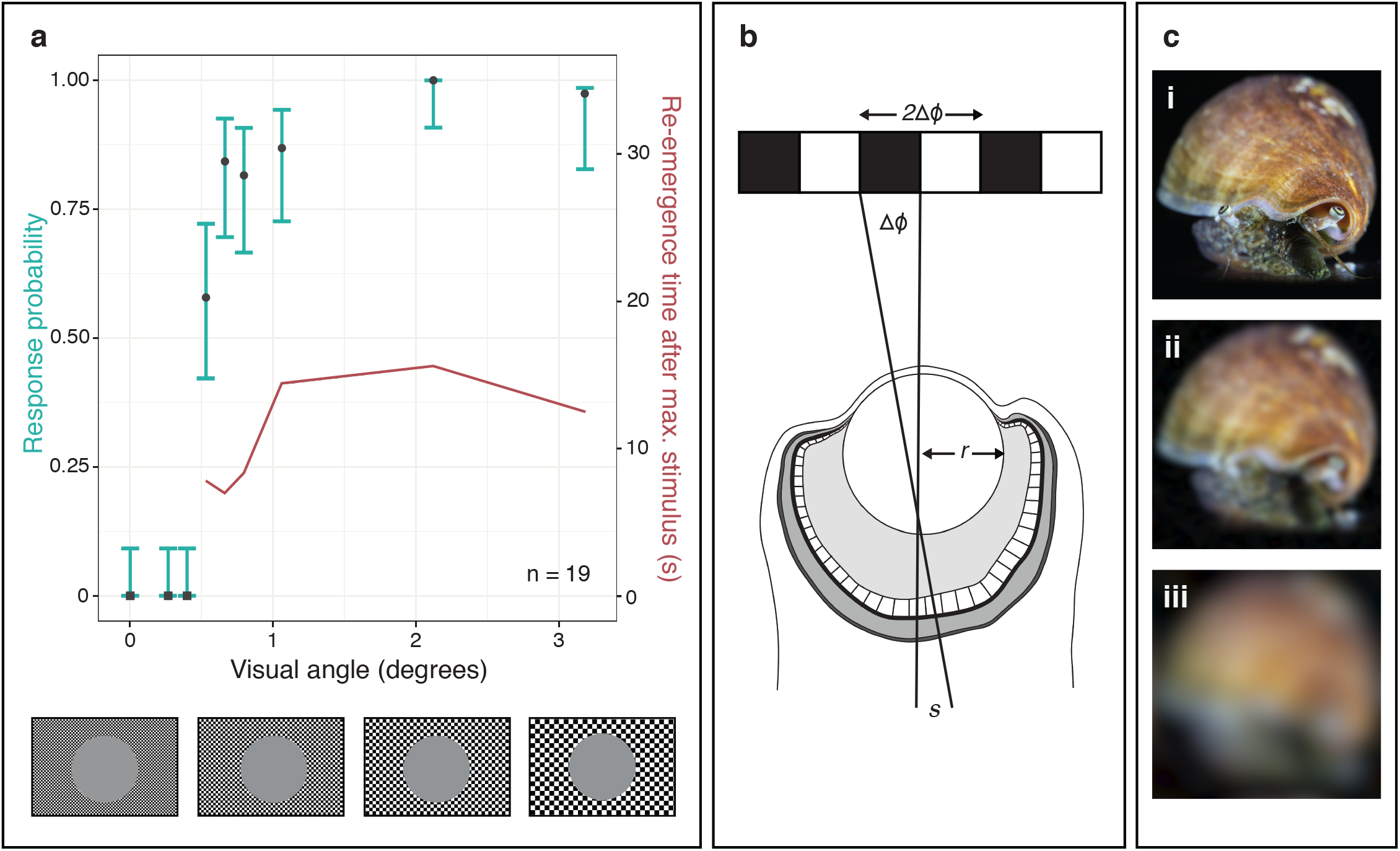
Calculation of visual acuity of *Conomurex luhuanus* based on (a) behaviour and (b) morphological data, with (c) depiction of acuity. (a) Response probabilities (black shapes) of *C. luhuanus* to expanding stimuli with visual angle of checkerboard background squares varied (below, not to scale): square, p = 1; circle, p = <0.01 following Bonferroni correction (n = 19). Error bars (green) are Wilson score intervals. Re-emergence time (red) is the time taken post-stimulus for ommatophores to return to their extended position and normal grazing to resume. This experiment included two replicates for every animal used. (b) Diagram illustrating the finest checkerboard that the eye can resolve, with an angular period of twice the inter-receptor angle Δφ. (c) Image of *Conomurex luhuanus* blurred using R v.4.0.3 (R Core Team, 2020) via the package AcuityView (Caves and Johnsen, 2017), according to the spatial resolution of: i) octopus (Land, 1981), ii) *C. luhuanus* (this study), iii) periwinkle *Littorina littorea* (Seyer, 1992).

### 3.2 SBF-SEM

#### 3.2.1 Anatomical studies suggest fine spatial resolution and high sensitivity

In this section, values are expressed as: mean ± s.d, standard deviation (s.e.m, standard error of the mean; n, number of measurements taken from the single *C. luhuanus* eye used for SBF-SEM). The spatial resolution estimated from anatomical data (*s*, 6.5 ± 0.9 mm (s.e.m 0.1 mm, n = 56); *f*, 720 mm (n = 1)) was calculated as 1.04 ± 0.14° (s.e.m 0.02°), twice the interceptor angle of 0.52 ± 0.07° (s.e.m 0.01°) (Fig. 4b; Eqn 1). The sensitivity value *S* of the *C. luhuanus* eye was calculated to be 7.78 ± 0.80 mm^2^.sr (s.e.m 0.05 mm^2^.sr) using anatomical data (*A*, 630 (n = 1); *f*, 720 mm (n = 1); *d*, 6.6 mm ± 0.8 (s.e.m 0.1 mm, n = 68); *x*, 70.9 ± 2.7 mm (s.e.m 0.6 mm, n = 20) (Eqn 2).

#### 3.2.2 Anatomical studies indicate at least six retinal cell types

The large *C. luhuanus* eye (diameter 1.2 mm, not including eyestalk tissue) contains a retina consisting of several layers. The retinal layer nearest to the vitreous body predominantly consists of the distal segments of the main photoreceptors. Dark pigment granules (ca. 0.5 mm) are found within most retinal cell types, predominantly within the pigmented region at the base of the distal segments (Fig. 5). Below this region, the main cell bodies form the nuclear layer, while their neuronal processes extend out and form a layer of neuropile nearest to the collagenous capsule, which is innervated by the optic nerve (Fig. 5).

**Figure 5.**
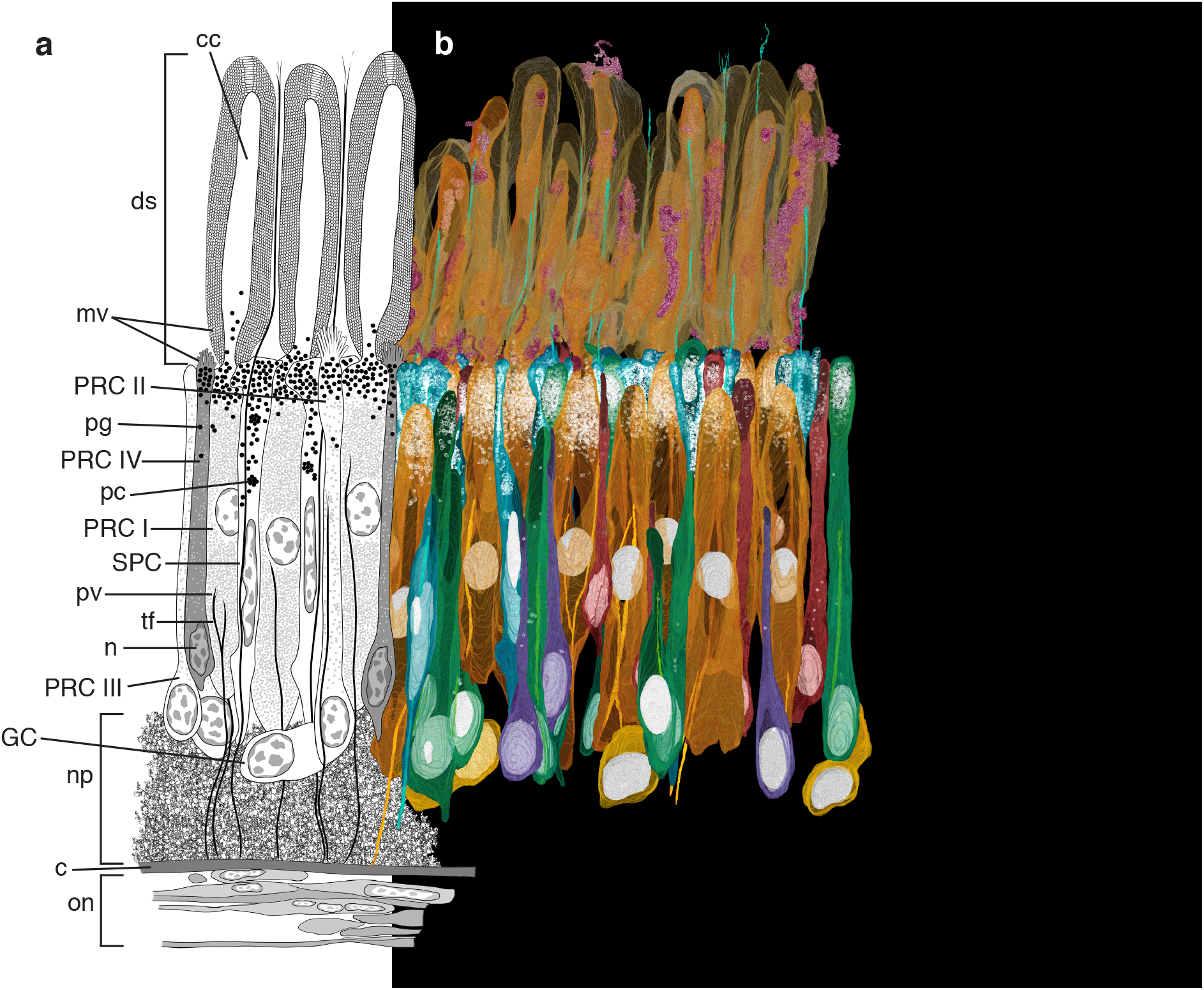
Retina structure of *Conomurex luhuanus*. a) Diagram (cell bodies and nuclei to scale) representing the supportive cell (SPC), ganglion cell (GC), and photoreceptor cells (PRC I-IV). b) Segmentation and reconstruction of cells using SBF-SEM data: blue, SPC; orange, PRC I; green, PRC II; purple, PRC III; red, PRC IV; yellow, ganglion cell; pink, phagocytic activity. Nuclei and pigment are highlighted in white. Abbreviations: c, capsule; cc, cytoplasmic core (of PRC I distal segment); ds, distal segments; mv, microvilli; n, nucleus; np, neuropile; on, optic nerve; pc, pigment cluster; pg, pigment granule; pv, photic vesicles; tf, tonofilaments. Scale bar = 20 mm. Note that following reconstruction the individual cells in b) have been layered consecutively using Adobe Photoshop v. 22.5.3 (Adobe Inc.) so that they are in the right position and overlying or underlying the correct adjacent cells. In this way, the semi-transparency of the cells, used in order to display pigment and nuclei, does not interfere with the clarity of the image (for unedited image, see Fig. S1). To ensure the image remained representative of the data, the positioning of the cells was verified using an underlying layer of the whole reconstructed retina for reference, since removed. All data was taken from a single eye of one specimen.

The area *C. luhuanus* retina sectioned by SBF-SEM data (2.048 × 10**-**_3_ mm^2^) contained 189 cells in the nuclear layer, from which an estimate of 1.57 × 10^5^ total retinal cells per eye was produced (Table 2). Cell types were distinguishable by differences in outer and inner segment morphology, electron density of the inner segment, and nucleus morphology and position, and various other cellular inclusions as described in the following sections. With SBF-SEM data, we could readily discern six morphologically distinct retinal cells, most clearly separable in the nuclear layer: a supportive cell (SPC), four photoreceptor cell types (PRC I-IV) and a ganglion cell (GC) (Figs 5, 6; Table 2). Cell types PRC IV and GC are newly described within the *C. luhuanus* retina from this study.

**Table 2.** Comparison of supportive cell (SPC), ganglion cell (GC) and photoreceptor cell types (PRC I-IV) within the *Conomurex luhuanus* retina.

| Cell type | Number of cells in $2.048 \times 10^{-3} \text{ mm}^2$ area sectioned (% total) | Mean cell volume in nuclear layer, $\text{mm}^3$ | Mean volume of pigment per cell, $\text{mm}^3$ | Mean pigment density, % of cell volume | Estimated cell count per $\text{mm}^2$ | Estimated cell count per eye |
| --- | --- | --- | --- | --- | --- | --- |
| SPC | 67 (35.4) | 116.7 | 10.4 | 8.9 | 32,700 | 55,600 |
| PRC I | 48 (25.4) | 565.2 | 14.8 | 2.6 | 23,400 | 39,800 |
| PRC II | 33 (17.5) | 214.3 | 4.9 | 2.3 | 16,100 | 27,400 |
| PRC III | 12 (6.3) | 120.0 | 0.11 | 0.09 | 5,900 | 10,000 |
| PRC IV | 21 (11.1) | 100.0 | 1.1 | 1.1 | 10,300 | 17,400 |
| GC | 8 (4.2) | 144.0 | 0 | 0 | 3,900 | 6,600 |
| **Total** | 189 | n/a | n/a | n/a | 92,300 | 156,900 |

**Figure 6.**
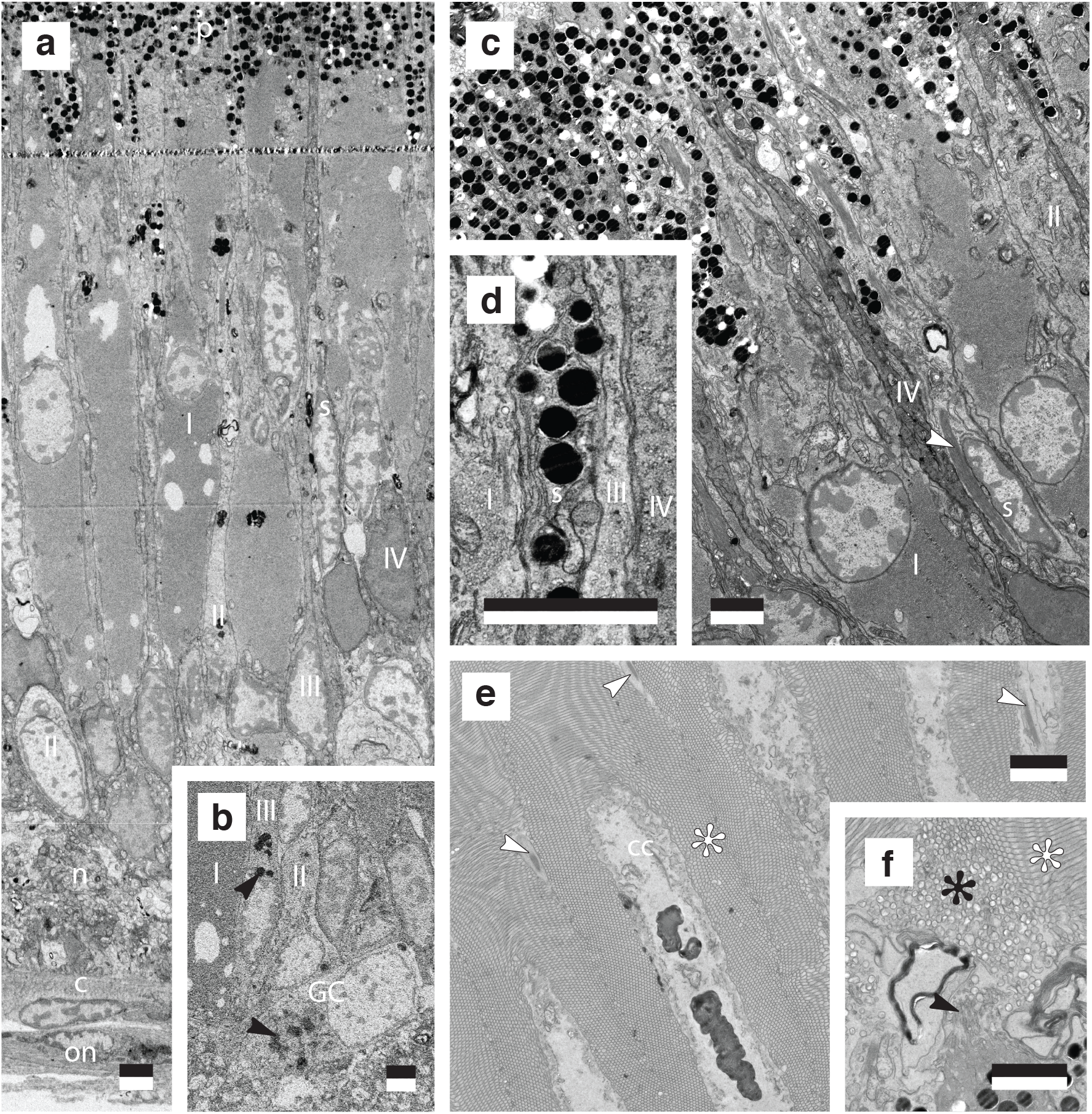
Structural characterisation of *Conomurex luhuanus* supportive (SPC) and photoreceptor retinal cells (PRC I-IV). TEM (a, c-f) and SBF-SEM (b) images show: a) cells within the nuclear region; b) cells close to the neural layer, with GC and PRC III containing large dense bodies (black arrow); c) cells at the nuclear and pigmented regions; d) differences in photic vesicle abundance among cell types; e) distal segments of PRC I; f) microvilli of PRC IV (black arrow) and PRC II (black asterisk). Abbreviations: c, capsule; cc, cytoplasmic core (of PRC I distal segment); n, neuropile; on, optic nerve; p, pigmented region; s, SPC; I-IV, PRC I-IV. White arrow: tonofilament; white asterisk: PRC II microvilli. Scale bar: 2.5 mm. All data was taken from a single eye of one specimen.

As sectioning did not occur precisely down the sagittal plane of the eye, photoreceptors with complete outer and inner sections could not be entirely traced. Therefore, photoreceptors were traced as near to completion as possible (Fig 5b; see Movie 2 for reconstruction images of each cell type), and a diagram produced based on these findings to illustrate the structures of each photoreceptor type, described below (Fig. 5a).

#### 3.2.3 Retina cell types

##### 3.2.3.1 Supportive cell (SPC)

Of the six cell types observed in the *C. luhuanus* retina (Figs 5, 6a-c), supportive cell (SPC) types are the most abundant (35.4% of total cells; Table 2). These cells are electron-lucent and lack photic vesicles (Table 2; Figs 5, 6a,c,d). The nuclei are characteristically narrow, elongated, and vary with respect to their positions within the nuclear layer (Figs 5, 6a,c; Movie 2). The cell body is also narrow, except within the pigmented region, where it expands to surround adjacent photoreceptor cells (Fig. 5; Movie 2). SPC is the most heavily pigmented retinal cell type (8.9% of the total cell volume; Table 2); unlike the other cell types, pigment granules extend from the pigmented region of the retina to as far as the nucleus (Figs 5, 6a,c,d; Movie 2). SPC pigment granules sometimes form clusters within the nuclear layer, bound by a membrane containing dense cytoplasm (Figs 5, 6d; Movie 2). A bundle of densely packed tonofilaments extends from the capsule (Fig. 7a), through the nuclear layer and between the photoreceptor cell type I distal segments in the rhabdomeric layer (Fig. 6c,e), finally dispersing in the vitreous body, or else prematurely between the distal segments (Fig. 7b,c). Within the rhabdomeric layer of the retina, filaments are surrounded by cytoplasm and a plasma membrane, which is often tightly packed around the filaments, though occasionally the cytoplasm and plasma membrane expand to occupy larger spaces between distal segments (Fig. 7d).

**Figure 7.**
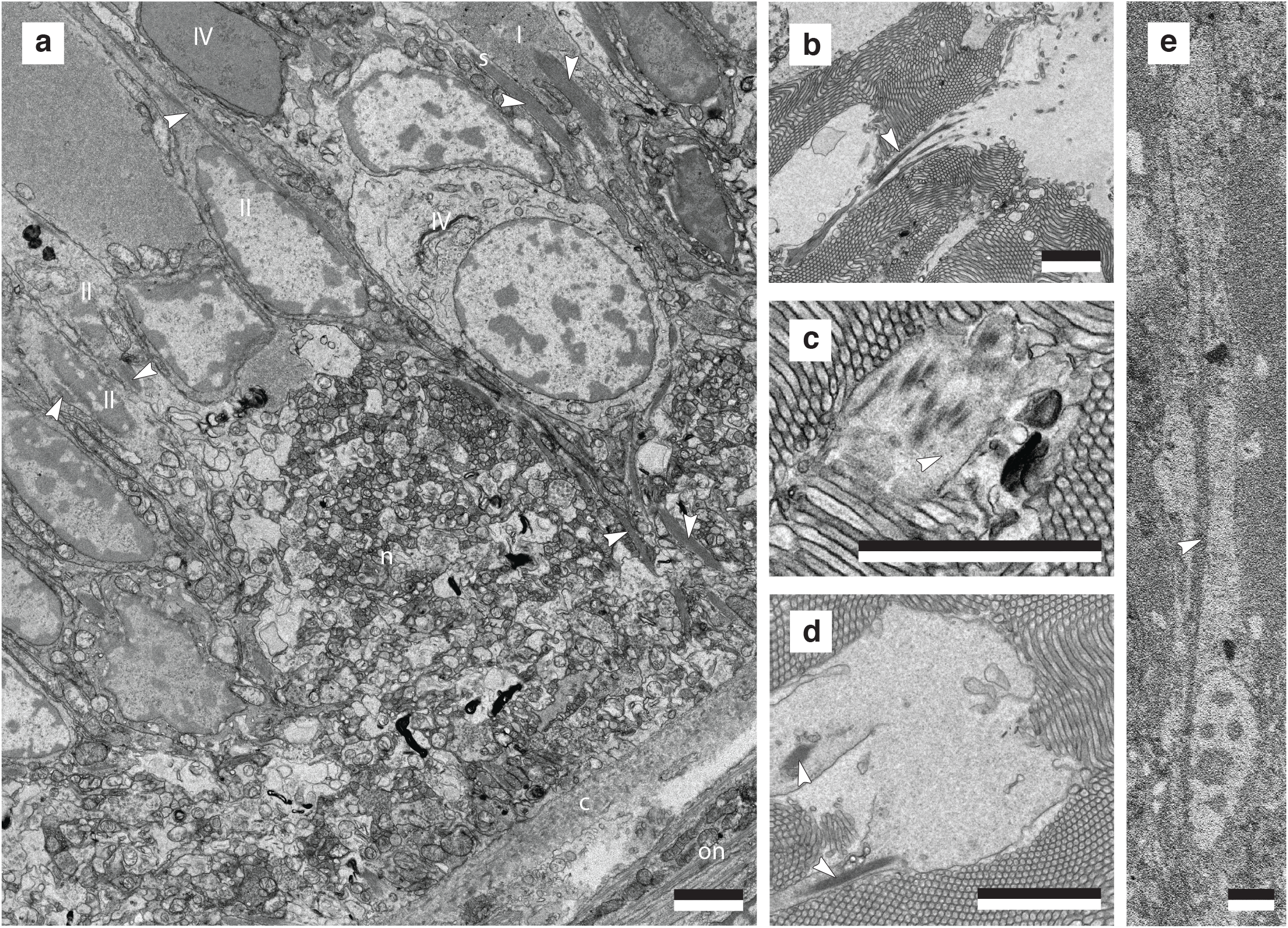
Morphological characterisation of filaments (white arrows) within *Conomurex luhuanus* retinal cells. TEM images show: a) bundles of filaments in supportive cells (SPC) and photoreceptor cells (PRC I-II) within the nuclear and neuropile layers; b) SPC tonofilament dispersing into vitreous body; c) SPC tonofilaments dispersing within the distal segment layer; d) cytoplasm and membrane not always tightly bound around SPC tonofilament. e) SBF-SEM image shows PRC II filaments extending midway through the nuclear layer before dispersing. Abbreviations: c, capsule; n, neuropile; on, optic nerve; s, SPC; I-IV, PRC I-IV. Scale bar: 2.5 mm. All data was taken from a single eye of one specimen.

##### 3.2.3.2 Photoreceptor cell type I (PRC I)

The main (most abundant; Table 2) photoreceptor cell, PRC I, possesses long (mean 70.9 ± 2.7 mm) distal segments that together comprise the majority of the rhabdomeric layer (Figs 5, 6d; Movie 2). The cell body of PRC I in the nuclear layer is wide, only tapering in the pigmented region and towards the neuropile, and contains ovoid nuclei positioned midway through the nuclear layer (Figs 5, 6a-c; Movie 2). The cytoplasm is packed with spherical vesicles, identified as photic vesicles (Fig. 6d), also containing bundles of filaments extending from the basal end of the retina to midway through the nuclear layer (Figs 5, 7a; Movie 2), though these filament bundles were rarely identified in the cell. In the distal segment layer, each long photosensitive organelle consists of a central cytoplasmic shaft extending out from the cell body, with an array of microvilli projecting from the surface, curving around the central shaft (Figs 5, 6d; Movie 2).

##### 3.2.3.3 Photoreceptor cell type II (PRC II)

PRC II is less abundant in the retina than SPC and PRC I, and has many short (7.1 ± 0.9 mm (s.e.m 0.2 mm, n = 20)) microvilli extending from the apical end of the cell body into the outer segment layer, instead of from a cytoplasmic core as in PRC I (Figs 5a, 6e; Movie 2). These microvilli are more electron-lucent and disordered compared to the regular arrangement of microvilli in the longer PRC I distal segments (Fig. 6f). Owing to the plane in which the retina was sectioned, microvilli in these SBF-SEM data were not able to be accurately reconstructed. The cytoplasm is electron-lucent and contains sparse photic vesicles, with a subspherical nucleus close to the neuropile (Figs 5–7; Movie 2). The cytoplasm contains bundles of filaments which extend from the neuropile to midway through the nuclear layer (Figs 5, 7a,e; Movie 2).

##### 3.2.3.4 Photoreceptor cell type III (PRC III)

PRC III is found infrequently within the retina and possesses a very narrow soma with electron-lucent cytoplasm, and a nucleus located close to the neuropile (Table 2; Figs 5–7; Movie 2). TEM images show sparse photic vesicles scattered in the cytoplasm (Fig. 6d), with large dense bodies identified in SBF-SEM sections (Fig. 6b). SBF-SEM data also revealed a lack of photopigment in the pigmented region, with sparse granules scattered in the nuclear layer (Table 2; Fig. 5; Movie 2).

##### 3.2.3.5 Photoreceptor cell type IV (PRC IV)

This cell type has not previously been observed in morphological studies of strombid retinas. PRC IV possesses very few microvilli projecting into the distal segment layer from a flat, apical surface (Figs 5, 6f). These microvilli are much shorter than those of PRC II-III (2.9 ± 0.3 mm (s.e.m 0.1 mm, n = 12)), and, like the remainder of the cell cytoplasm, are very electron dense compared to all other cell types (Figs 6, 7a). The cell body is of a similar overall shape to that of PRC II: narrow in the pigmented region and much of the nuclear layer, widening at its irregularly-shaped nucleus near the neuropile (Figs 5–7; Movie 2).

##### 3.2.3.6 Ganglion cell (GC)

Like PRC IV, GCs have not previously been identified in strombid retinas and is the least frequent component of the retina (Table 2). This ganglion cell is variable in shape and size with a large, ovoid nucleus (Figs 5, 6b). The cytoplasm is electron lucent and lacks photic vesicles, instead containing numerous large dense bodies (Fig. 6b).

#### 3.2.4 Phagosomes

SBF-SEM and TEM data were also used to identify electron-dense, lamellar inclusions in the distal segment layer of the retina, identified as phagosomes (Figs 5b, 8). Their structures, albeit diverse, are all comprised of concentric systems of membranes, mostly irregular in shape, with few circular types (e.g. Fig. 8a). They have no specific intracellular location within the cytoplasmic core and can be found in the basal or apical portions of the cell (Figs 5; 8b-e), with others located within vacuole-like structures just above the apical end of the distal segments in the vitreous body (Fig. 8a; Movie 2). Other phagosomes are located between the main distal segments, surrounding the microvilli projecting from the accessory photoreceptor cells (Fig. 5b; Movie 2).

**Figure 8.**
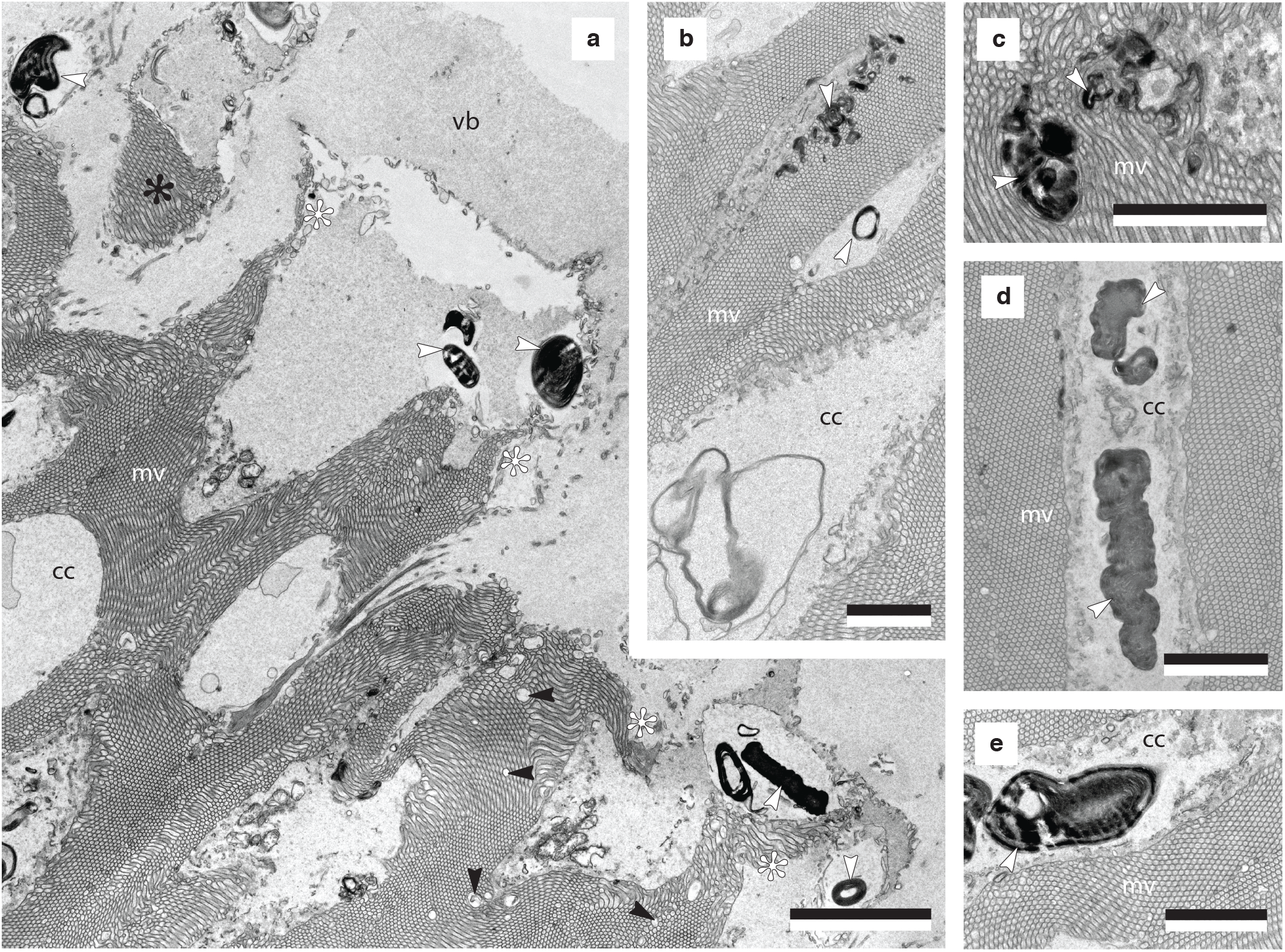
Morphological characterisation of phagosomes (white arrows) within the main *Conomurex luhuanus* photoreceptor cell (PRC I) distal segment layer. TEM images show: a) phagosomes at apical end of distal segments, with punctuate discontinuities in the microvillar array (black arrow), and apical ends of distal segments curling away (white asterisk) or completely separated (black asterisk) from rest of distal segments; b,c) phagosomes intruding on microvillar arrays; d,e) phagosomes within cytoplasmic core of distal segments. Abbreviations: cc, cytoplasmic core; mv, microvilli; vb, vitreous body. Scale bar: 2.5 mm. All data was taken from a single eye of one specimen.

## 4 Discussion

This is the first study to link visually-influenced behaviours in conch snails to the structure and function of their well-developed eyes. In this study, the use of behavioural experiments together with volumetric electron microscopy has enabled new insights into the strombid visual system, the findings of which are discussed below.

### 4.1 Strombids have high spatial resolution

Estimates of spatial resolution from behavioural (αmin 1.06°) and anatomical (2Δ*ϕ* 1.04 ± 0.14°) measurements both suggest that the strombid eye can resolve a objects of ca. 1° in the visual field (Fig. 4). These estimates of visual acuity from the anatomy of the *C. luhuanus* eye mirror previous estimates from another strombid species, *Lobatus raninus* (2Δ*ϕ* 0.94°; Seyer, 1994). However, while the potential for high resolution strombid vision was suggested by Seyer (1994), the combined behavioural and anatomical approaches used in this study provide conclusive evidence that these animals utilize this resolution in visual tasks. A spatial resolution of 1.0° in the strombids is much finer than is estimated for other gastropods; these vary from 3.6° in *Littorina littorea* (anatomical data; Seyer, 1992), to 52° in *Arion rufus* (behavioural data; Zieger et al., 2009). Like the strombids, these are mostly non-predatory gastropods. Apart from the predatory heteropods (Land, 1982), the only other molluscs that exhibit similar resolutions to conch snails are predatory cephalopods which use high-resolution camera type eyes to target prey; these range widely in resolution, from 0.57° in cuttlefish *Sepia officinalis* (behavioural data; Groeger et al., 2005) and 0.50° in squid *Japetella* (anatomical data; Sweeney et al., 2007), to 0.04° in *Octopus vulgaris* (anatomical data; Young, 1962a; Land, 1981).

The ability of *C. luhuanus* to utilize high resolution information is also indicated by the high density of main photoreceptor cells in its retina (PRC I), estimated to be 3.98 × 10^4^ per eye (Table 2), though this does not consider the possible variation in cell density across the retina, as noted for cephalopods (e.g. Young, 1962a). This estimate of PRC I cells in *C. luhuanus* is less than the 5 × 10^4^ PRC I cells suggested by Gillary and Gillary (1979), and closely matches the 4×10^4^ optic nerve fibres estimated from the same study. Nevertheless, both estimates of PRC I suggest more numerous photoreceptors are present in the eyes of conch than in other gastropods, such *Helix* and *Onchidium*, with 2500–3800 and 600 main photoreceptor cells per eye, respectively (Brandenburger, 1975; Katagiri et al., 1995); by comparison, *Octopus vulgaris* is estimated to possess ca. 2.0 ×10^7^ photoreceptors per eye (Young, 1962a). Hess (1905) estimates the photoreceptor density in *Sepia officinalis* to be 105,000 per mm^2^; however, in *O. vulgaris*, the density (70,000 per mm^2^) is not dissimilar to that within *C. luhuanus* (55,700 per mm^2^ PRC I-IV, or 23,400 per mm^2^ PRC I only), although in *O. vulgaris*, 25% of the cells in the nuclear layer are supporting cells, as opposed to 37% in *C. luhuanus* (excluding ganglion cells; Table 2; Young, 1962a).

Spatial resolution is one of several variables which determine the visual tasks an eye can support. In particular, it is an important property for detecting moving objects in the environment, especially potential predators (Nilsson, 2013). Like other strombid species, *C. luhuanus* are commonly found in large aggregations of 100–200 individuals, between ca. 0.5 and 30 individuals per m^2^; an easy visual target for some predators (Poiner and Catterall, 1988; Ulm et al., 2019). While chemosensory perception and their subsequent leaping escape behaviour (facilitated by long opercula) help strombids avoid slow-moving predators such as cone snails (Berg 1974, 1975; Field, 1977), their escape response is too slow to be effective against fast-moving predators such as fish, shell-peeling crabs, or octopuses (Savazzi, 1991). Facing these quicker predators, strombids must detect approaching objects and withdraw into their shell, relying on its passive mechanical protection (Savazzi, 1991), though the shell can be crushed or bored through by certain predators (Berg, 1974).

The behavioural experiments demonstrated that an expanding circle subtending 2.3–9.8° of the visual field was capable of causing *C. luhuanus* to stop feeding and slowly withdraw their eyestalks and proboscis. However, additional defensive responses only occurred after the stimulus expanded to an angular size of 11.8° (Fig. 2a). These different behaviours indicate a possibility of two behavioural thresholds in response to approaching objects: firstly, the point at which the snail ceases other activity to focus on the approaching object, and secondly, the point at which the snail actively avoids a predator. This indicates that one function of high visual acuity in strombids is likely for detection of potential predators as early as possible. Behavioural studies of other non-predatory molluscs with poorer spatial resolution have demonstrated a variety of uses of visual information: similarly detecting potential predators [e.g. bivalve *Cardium edule* (Barber and Wright, 1969) and scallop *Argopecten irradians*, at a long range (Chappell et al., 2021)], orienting to celestial cues and finding suitable habitats (e.g. *Littorina*; Fig. 4c; Newell, 1958; Hamilton, 1977). Therefore, previous authors have suggested that vision in strombids may also support other behavioural tasks, such as to help maintain a straight path upon escaping molluscivorous cone snails (Field, 1977), to strike a predator more accurately with a kick of its long, serrated operculum as a deterrent (Prince, 1955; but see Berg, 1974), or to find conspecifics and suitable habitats. However, these are visually guided behaviours yet to be tested.

### 4.2 Strombids have both high contrast sensitivity and absolute sensitivity

In addition to high spatial resolution, high contrast sensitivity is also advantageous for early predator detection. Experiments with the expanding visual stimuli showed *C. luhuanus* to be capable of discriminating between small differences in light intensity (Michelson contrast 0.07; Fig. 3). The ability to detect small changes in contrast within any environment allows for a larger, safer distance at which potential predators are identified (Land, 1981; Smolka and Hemmi, 2009).

With regard to the absolute sensitivity of the eye, increased photon capture is facilitated through several adaptations seen in strombid eyes (Hughes, 1976; Gillary and Gillary, 1979; Seyer, 1994), including wide apertures to allow more light into the eye, long distal segments of the photoreceptors to increase the absorbance path length, and photoreceptors with wide acceptance angles (Land and Nilsson, 2012). The absolute sensitivity value, *S*, of the *C. luhuanus* eye (7.78 ± 0.80 mm^2^.sr), is less than values previously calculated for the strombid *Lobatus raninus* (9.9 mm^2^.sr; Seyer, 1994); differences may be accounted for by the large variation in rhabdom length in the retina (Gillary, 1974, pl. 1; *pers. obs*.), meaning that measurements taken from the area sectioned in this study may not be representative of the sensitivity of the whole eye. These values are nevertheless similar to those calculated for octopus (9.7 mm^2^.sr; Hanlon and Messenger, 2018), which often inhabit similar coastal sea floor environments to *C. luhuanus* (predominantly sand, coral rubble and seagrass beds; Poiner and Catterall, 1988; Ulm et al., 2019).

Sensitivity estimates from morphological data may also reflect the fact that strombids appear to be most active in dim light [*Lobatus raninus* (Seyer 1994) and *C. luhuanus* (TMC *pers. comm*.; *pers. obs*.)], though laboratory and field studies of strombids have observed some feeding activity around the clock (Randall, 1964; *pers. obs*.). Comparisons between the eyes of known nocturnal and diurnal species suggest that strombid *S* values are congruous with twilight activity (Seyer, 1994; Land and Nilsson, 2012), a predication that is further supported by the existence of several high sensitivity adaptations of the strombid eye. These include the long length of the PCR I distal segments (mean 70.9 ± 2.7 mm; Fig. 5) and their specialised structure: a cytoplasmic core extending from the pigmented region of the cell, around which arrays of microvilli are circularly arranged (Fig. 6d; Hughes, 1976; Gillary and Gillary, 1979). The cytoplasmic core is suggested as an adaptation for more efficient transport of materials in longer distal segments (Hughes, 1976); however, this structure may also increase sensitivity due to microvillar orientation. In both the circular arrays of microvilli in strombids and the brush-like arrays in gastropods *Bulla, Limax, Deroceras* and *Athoracophorus* (Katoaka, 1975; Eakin et al., 1980; Jacklet and Colquhoun, 1983; Newell and Newell, 1968), microvilli are oriented perpendicular to the incident light entering the eye, allowing maximum absorption of photons and thereby increasing visual sensitivity (Eakin et al., 1980). This is consistent with the fact that *Bulla, Limax, Deroceras* and *Athoracophorus* are nocturnal (Carmichael, 1931; Stephenson, 1968; Eakin et al., 1980; Jacklet and Colquhoun, 1983), and that strombids are also observed to be active in dim light.

### 4.3 The strombid retina is composed of six different cell types

If a measure for complexity is the number of different parts of which a given structure is composed (McShea, 2000; Oakley and Rivera, 2008; Arendt et al., 2009), the six cell types within the strombid retina indicates a higher complexity than that of other gastropod retinas known from morphological studies. In contrast to the six retinal cells (four photoreceptors, 1 ganglion cell and one supportive cell) identified in *C. luhuanus* (Table 2; Figs 5–7; Movie 2), most gastropods possess only two or three retinal cell types: a supportive cell, a main photoreceptor cell, and sometimes an accessory photoreceptor cell with shorter distal segments (e.g. Eakin et al., 1967; Jacklet and Colquhoun, 1983; Seyer, 1992, 1998; Pinchuck and Hodgson, 2018). An exception to this is seen in the retinas of *Helix aspersa* and *Onchidium verruculatum*, wherein a fourth (Brandenburger, 1975) or fifth (including ganglion cell; Katagiri et al., 1995) cell type was identified but not described with detail due to low frequency in the retina. Despite belonging in the same superfamily as the strombid *C. luhuanus*, only two cell types are identified in aporrhaid *Aporrhais pespelecani* (Blumer, 1996). These comparisons indicate a remarkably complex visual system in *C. luhuanus* compared to most other gastropods.

Previous studies using TEM alone suggest that the *C. luhuanus* retina contains four different cell types: a supportive cell (SPC) and three types of photoreceptors (PRC I-III) (Gillary and Gillary, 1979; Ozaki et al., 1986). Histological studies in other strombid species identified only two of these cells, SPC and PRC I (Prince, 1955; Hughes, 1976). Of the six cell types identified in this study using SBF-SEM and TEM data, two are newly identified (PRC IV and GC), and four matched those described in *C. luhuanus* in previous studies (Gillary and Gillary 1979; Ozaki et al., 1986; Table 2). However, PRC III is only putatively identified due to the limited description in Gillary and Gillary (1979) and differences in imaging resolution between data in this and previous studies. Nevertheless, the possible fourth cell described by Gillary and Gillary (1979) and PRC III share a very narrow soma, electron-lucent cytoplasm, infrequent occurrence, and position of the nucleus close to the neuropile, indicating a strong likelihood that these are the same cell type (Table 2; Figs 5–7; Movie 2). Furthermore, SBF-SEM data did not identify microvilli at the apical end of PRC III, also not described in previous work (Gillary and Gillary, 1979; Ozaki et al., 1986).

Despite several differences, these retinal cells share key features with those reported for other gastropod species, allowing discussion of the likely function of the cell types described in this study (Figs 5–7). Unlike supportive cells, gastropod photoreceptor cells contain many small electron-lucent cytoplasmic vesicles, referred to as photic vesicles (e.g. Eakin et al., 1967; Eakin and Brandenburger, 1975; Stoll, 1973; Hughes, 1976; Gillary and Gillary, 1979; Gibson, 1984; Eakin, 1990; Pinchuck and Hodgson, 2018). The distribution of these vesicles varies across groups and between cell types; in cells such as PRC I in *C. luhuanus*, photic vesicles are densely packed throughout the cytoplasm (e.g. Pinchuck and Hodgson, 2018; Fig. 6d), whereas investigations within slugs revealed aggregations just beneath the light-sensitive microvilli in the light-tolerant *Ariolimax* or concentrated basally near the nuclei in the nocturnal *Limax*, supporting the suggestion that the vesicles are associated with photoreception (Eakin and Brandenburger, 1975). Previous studies suggest several functions for these vesicles, including storage of photopigment (Röhlich and Torok, 1963; Eakin and Brandenburger, 1968; Eakin, 1990). Within *C. luhuanus* (Ozaki et al., 1986) and *Onchidium* (Katagiri et al., 2001), an abundance of the photopigment retinochrome was found in fractions of photoreceptor cells containing photic vesicles, supporting the idea that these are involved in storage. The presence of these vesicles in PRC I-IV within *C. luhuanus* (albeit more sparsely in PRC II-IV; Gillary and Gillary, 1979; Fig. 6d) therefore indicates that these cells are involved in photoreception.

Unlike the photoreceptor cells, the newly described ganglion cell in this study lacks photic vesicles, instead containing numerous large dense bodies, likely secondary residual lysosomes as identified in other gastropod studies (Fig. 6b; Eakin et al., 1980). This cell is analogous to a variety of cells located exclusively in the neural layer of the retina in previous gastropod studies, described as ganglion cells, secondary cells, and neurosecretory cells or neurons (Figs 5, 6b; Stoll, 1973; Brandenburger, 1975; Eakin et al., 1980; Jacklet and Colquhoun, 1983; Katagiri et al., 1995). The newly described cell PRC IV differs predominantly from other retinal cells in that its cytoplasm is very electron-dense and contains no bundles of filaments (unlike SPC and PRC I-II; Fig. 7). Furthermore, the euchromatin and heterochromatin within the nucleus of PRC IV is much more electron-dense than in the other retinal cells (Figs 6a,c, 7a), more closely resembling that of the dense photoreceptor in the gastropod *Bulla* (Jacklet and Colquhoun, 1983). A similar photoreceptor cell was observed in the eye of *Ilyanassa*, which shows the same dense cytoplasm, narrowing of the cell body towards the pigmented region and irregular, small microvilli as PRC IV, though, unlike PRC IV, lacks electron-lucent photic vesicles (Gibson, 1984, figure 8; Movie 2; Figs 5, 6f).

Though some clues as to cellular function are given by structural features as discussed above, further work is required to investigate what implications there are for the four strombid photoreceptors on visual processing. Electrophysiological investigations into the neural mechanisms of the *C. luhuanus* visual system indicated that photic stimulation triggers highly complex neural interactions, involving excitation, inhibition, and oscillatory ‘off’ activity, unlike gastropods *Otala* and *Helix* which exhibited only excitation activity (Goldman and Hermann, 1967; Gillary, 1970, 1974, 1977). Gillary (1974) suggested that some processing of neural information occurs in the retina, similar to vertebrate retinae which act as complex filters to transfer specific information (including motion, contrast, colour and resolution) about images to the brain in parallel via different classes of ganglion cells (Wässle, 2004; Knudsen, 2020). By contrast, cephalopods do not possess ganglion cells (Yamamoto et al., 1965) and visual information processing is solely undertaken in the large optic lobe of the brain (Young, 1962b; Williamson and Chrachri, 2004; Hanke and Kelber, 2020). Therefore, the strombid visual processing suggested by Gillary (1974) and supported by this study, as well as the diversity of photoreceptor cell types in the retina, suggests a different evolutionary path taken to the visual processing within cephalopods. The convergent evolution of large, high-acuity camera-type eyes in cephalopod and conch snails, yet non-convergent visual processing, makes the strombids an interesting subject for understanding the hierarchical steps in visual data processing and the evolution of vision in molluscs.

## 5 Conclusion

This study provides behavioural evidence that the strombid gastropod *C. luhuanus* has high contrast sensitivity and high visual acuity, able to respond to a Michelson contrast of ca. 0.07 and differences as small as 1.06° in the visual field. The estimated spatial resolution from behavioural data is strongly supported by an estimate of 1.04 ± 0.14° from anatomical data. This is the most acute vision described for any non-predatory gastropod and supports previous estimates based on morphological data, demonstrating the value of integrating morphological and behavioural approaches. Withdrawal responses to expanding stimuli suggest that high visual acuity and sensitivity is likely to play a vital role in early predator detection in this species; however, this resolution seems far superior to that required for this task when compared to visual acuity in other gastropods, and it is probable that high spatial resolution also underpins other behaviours in strombids.

New techniques (SBF-SEM, in conjunction with TEM) reveal six kinds of retinal cells within the *C. luhuanus* retina: a supportive cell, a ganglion cell, and four photoreceptor type cells I-IV. Two of these cells, the ganglion cell and the fourth photoreceptor cell are newly discovered and described for the first time in this study: These data provide new insights into cell functions and widens our understanding of the complexity of the retina structure in strombids. These findings suggest that strombids have a remarkably complex retina compared to those within cephalopods and other gastropod groups, suggesting differences in the way visual information is processed among molluscs.

## Acknowledgments

We are grateful to Chris Neal, Sally Hobson, and the Wolfson Bioimaging Facility (University of Bristol) for carrying out specimen embedding, mounting and SBF-SEM imaging, in addition to initial TEM imaging prior to sectioning. We thank James Chen, Martin How and the Animal Services Unit (University of Bristol) for tank maintenance and care of the conch snails. We also thank Michael Bok (Lund University) for providing an image of one of the animals used in this study, used in Fig. 4c for the purpose of illustrating differences in spatial resolution, and in Movies 1 and 2. We are grateful to Vincent Fernandez and Brett Clark (Natural History Museum, London) for facilitating remote access to VGStudio Max software.

## Competing interests

No competing interests declared.

## Funding

This paper is supported by the NERC GW4+ Doctoral Training Partnership [grant reference NE/L002434/1].

## Notes

### Competing Interest Statement

The authors have declared no competing interest.

